# Autophagy-mediated CTR1 turnover orchestrates the reciprocal interaction between autophagy and ethylene signaling

**DOI:** 10.1101/2024.07.31.606019

**Authors:** Hye Lin Park, Weiwei Zhang, Yuan-Chi Chien, Chanung Park, Gyeong Mee Yoon

## Abstract

The phytohormone ethylene and autophagy are crucial for plant adaptation to various environmental stresses, yet the integration of these signaling networks into stress responses is not fully understood. Here, we report that ethylene signaling and autophagy reciprocally regulate each other through Constitutive Triple Response 1 (CTR1), a negative regulator of ethylene signaling. Autophagy facilitates the turnover of the CTR1 protein, which interacts with the key autophagy-related protein ATG8 as autophagic cargo. Impaired autophagy attenuates ethylene responses. Conversely, ethylene-insensitive mutants exhibit enhanced autophagic flux, while a constitutive ethylene response mutant is hypersensitive to carbon starvation stress, which induces autophagy. This suggests that ethylene suppresses autophagy during carbon limitation. We further elucidated that ethylene receptors with a receiver domain play a primary role in modulating autophagy, while receptor kinase activity is not essential. Our findings uncover that the autophagic control of CTR1 degradation allows reciprocal cross-regulation between autophagy and ethylene transduction cascades, optimizing stress responses and resilience.

## Main text

When stressed, plants reprogram their cellular, biochemical, and physiological features to acclimate to prevailing stress conditions. During this process, it is critical to keep their stress responses within optimal threshold levels—strong enough to withstand yet restrained enough to recover when the stress subsides. Ethylene signaling and autophagy, a self-eating cellular degradation and recycling process, are two well-known cellular pathways largely responsible for plant acclimation to a broad range of stresses^1–7^. However, despite well-established signaling pathways and physiological outputs, there is still a limited understanding of how these two pathways optimize the levels of stress responses to confer adequate stress acclimation and resilience.

Ethylene is a key plant hormone that enables plants to cope with environmental stresses. It also plays an important role in plant growth and development by regulating fruit ripening, seedling germination, senescence, and root hair formation^7^. Ethylene is perceived by ethylene receptors located on the endoplasmic reticulum (ER) membrane^8, 9^. At low levels of ethylene, these receptors activate Constitutive Triple Response 1 (CTR1), a Raf-like protein kinase, which phosphorylates Ethylene Insensitive 2 (EIN2), a key positive regulator of ethylene signaling. Phosphorylation of EIN2 by CTR1 leads to its inactivation, likely through degradation mediated by EIN2-targeting protein 1 (ETP1) and ETP2^10–14^. In the presence of ethylene, the ethylene receptors and CTR1 proteins are inactivated, which results in decreased phosphorylation of EIN2. This triggers the cleavage and transport of the C-terminal domain of EIN2 (EIN2-CEND) into the nucleus and processing body (P-body), where EIN2-CEND indirectly activates the EIN3 transcription factor and sequesters mRNA of EIN3-binding F-Box proteins (EBFs), respectively, to activate ethylene responses^15, 16^. Similarly, CTR1 is released from the ER to the cytoplasm upon ethylene binding to the receptors. CTR1 then translocates into the nucleus, where it enhances ethylene responses by stabilizing EIN3 proteins through direct interaction with EBFs^6^. The CTR1-EBF interactions presumably suppress EBF function, although the underlying mechanism is unknown.

The protein turnover of several ethylene signaling components, such as EIN2, EIN3/EIN3-like (EIL), and EBFs, is relatively well-established^12, 14, 17–19^. However, the mechanism governing CTR1 turnover remains elusive despite its critical role in the ethylene signaling pathway. Ethylene and 1-aminocyclopropane-1-carboxylic acid (ACC) have been shown to increase CTR1 protein levels in *Arabidopsis* plants, which facilitates the nuclear translocation of CTR1 and turns it into a positive regulator for the ethylene response^6, 20^. This suggests that cellular CTR1 levels, along with EIN2 and EIN3/EILs, play an important role in modulating plant stress responses. Although the mechanism underlying ethylene-induced stabilization of CTR1 is unknown, these results imply that ethylene negatively regulates the mechanism governing CTR1 protein turnover, thus contributing to optimal levels of ethylene responses in plants.

The diverse roles of ethylene in modulating various aspects of plant growth and mediating adaptive responses to biotic and abiotic stresses are underpinned by complex signaling crosstalk between ethylene and other cellular processes. One such process emerging as a key regulator of ethylene signaling is autophagy, a catabolic mechanism employing vacuolar degradation and recycling of cellular components to ensure plant survival under stress. Ethylene and autophagy regulate several key developmental transitions and stress response pathways, including senescence, hypoxia, drought tolerance, and nutritional stress adaptation^4, 5, 21–29^. This implies significant convergent signaling and physiological crosstalk between the two pathways. However, despite several potential nodes of interaction, the molecular mechanism underlying ethylene-autophagy crosstalk remains largely obscure.

In this study, we elucidate the reciprocal interaction between ethylene signaling and autophagy through the autophagic turnover of CTR1, revealing a crosstalk node integrating these pathways. Using autophagy-defective and ethylene signaling mutants, we demonstrate that autophagy positively regulates ethylene responses by destabilizing CTR1 proteins.

Conversely, ethylene negatively regulates autophagy by destabilizing autophagosomes and modulating the expression of autophagy-related (ATG) genes and Target of Rapamycin (TOR) signaling components under carbon starvation. Furthermore, ethylene receptors with a receiver domain are crucial for regulating autophagic responses independently of kinase activity.

Together, our work demonstrates how plants coordinate the control of CTR1 turnover to dynamically co-regulate the ethylene and autophagy pathways, optimizing stress responses and resilience to nutritional deficiencies while sustaining growth.

## RESULTS

### CTR1 protein levels are regulated by autophagy

CTR1 plays a critical role in modulating responsiveness to ethylene in multiple developmental and physiological responses^6, 30–33^. However, the mechanism regulating the level of CTR1 protein is unknown. Given potential links between autophagy and ethylene function, we examined whether autophagy plays a role in CTR1 protein turnover under carbon starvation, a well-known environmental condition for inducing autophagy^22, 34^. We transferred 7-day-old light-grown wild-type (WT) *Arabidopsis thaliana* seedlings to sucrose-deficient media and incubated them in darkness. Within 24 hours, the steady-state levels of CTR1 protein were substantially decreased and undetectable by 72 hours. However, treatment with concanamycin A (ConA), an inhibitor of vacuolar degradation, not only blocked CTR1 reduction but also led to gradual accumulation (**Fig. 1a**)^34^. Unlike in WT seedlings, genetic disruption of autophagy in the *atg7-2* mutant seedlings abolished the CTR1 decrease under these conditions (**Fig. 1a**)^35^. Notably, we observed that CTR1 protein levels also started to reduce after 4 days of dark incubation in the *atg7-2* seedlings, likely attributed to the intrinsic chlorosis phenotype of the *atg7-2* mutant.

**Figure 1.**
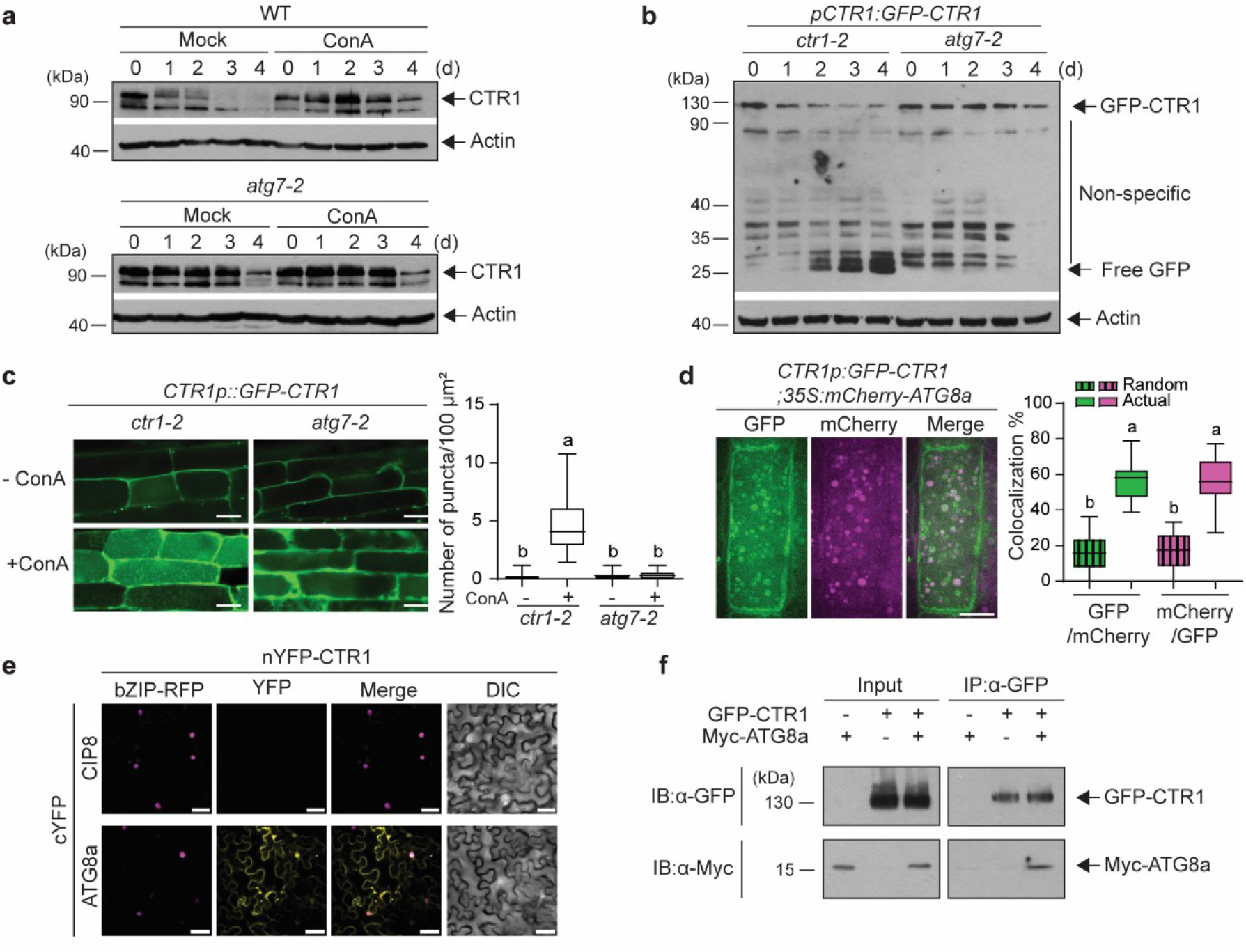
CTR1 is degraded by the autophagy pathway in *Arabidopsis*. **a**. Concanamycin A inhibits CTR1 degradation. Seven-day-old WT and *atg7-2* mutant seedlings grown in light were transferred to sucrose-deficient liquid media with or without concanamycin A (ConA, 0.5 μM). The seedlings were then incubated in darkness for the indicated times to induce carbon starvation. Total proteins were extracted and analyzed by immunoblotting using anti-CTR1 antibodies. Anti-actin antibodies were used as a loading control. **b**. Time course of free GFP release from GFP-CTR1 during carbon starvation. *ctr1-2* and *atg7-2* seedlings expressing GFP- CTR1 were carbon starved for the indicated times. Total protein extracts were immunoblotted with anti-GFP antibodies. Arrows indicate the GFP-CTR1 fusions and the free GFP released upon degradation. **c**. Delivery of GFP-CTR1 to vacuoles is blocked in *atg7-2*. Five-day-old light-grown WT and *atg7-2* seedlings expressing GFP-CTR1 were incubated in a sucrose-free medium with ConA under light or dark conditions for 16 hours. Root cells were examined for autophagic vesicles by confocal microscopy. Scale bars, 10 μm. One-way ANOVA with Tukey’s post-hoc test was performed, and the letters denote statistically significant differences between groups, *p* < 0.0001 (n ≥ 20 cells from five to six seedlings for each treatment) **d**. GFP-CTR1 colocalizes with mCherry-ATG8a. Seedlings co-expressing GFP-CTR1 and mCherry-ATG8a were incubated in a sucrose-free medium with ConA for 16 hours. Root cells were imaged by light sheet fluorescence microscopy to visualize the intracellular colocalization of GFP-CTR1 and mCherry-ATG8a. As a negative control, random colocalization was measured by rotating one image 180 degrees and repeating the analysis on the same image pair, assessing the expected colocalization due to chance. Scale Bar, 10 μm. One-way ANOVA with Tukey’s post-hoc test was performed, and the letters denote statistically significant differences between groups, *p* < 0.0001 (n ≥ 5 seedlings). Two different sections in the cells of each seedling were imaged and measured. **e**. CTR1 interacts with ATG8a in *Nicotiana benthamiana*. Leaves of *N. benthamiana* were co-infiltrated with agrobacteria carrying plasmids for nYFP-CTR1, cYFP- ATG8a, or cYFP-CIP8 (negative control), and bZIP-RFP (expression control). After 3 days, the reconstitution of YFP signals was observed by confocal microscopy, indicating CTR1-ATG8a interaction. **f**. Co-immunoprecipitation of CTR1 and ATG8a. Leaves of *N. benthamiana* were infiltrated with agrobacteria expressing GFP-CTR1 alone, myc-ATG8a alone or both GFP-CTR1 and myc-ATG8a. Total protein extracts were incubated with anti-GFP magnetic beads to pull down GFP-CTR1. The immunoprecipitants were subsequently immunoblotted with anti-GFP and anti-Myc antibodies to detect the co-immunoprecipitation of CTR1 and ATG8a.

Additionally, we utilized GFP-cleavage assays to evaluate the autophagic turnover of GFP-CTR1 in WT and *atg7-2* mutant (**Fig. 1b**)^34, 36, 37^. This assay determines the vacuolar delivery of GFP fusion proteins by quantifying the accumulation of protease-resistant free GFP in vacuoles.

Under carbon starvation, we observed escalating levels of liberated GFP from the GFP-CTR1 while descending levels of GFP-CTR1 in a *ctr1-2* mutant background. However, carbon starvation-triggered GFP release was absent in the *atg7-2* mutants (**Fig. 1b**). These results demonstrate that CTR1 protein levels are regulated by autophagy-mediated trafficking and subsequent vacuolar degradation under carbon deficit stress.

As CTR1 protein levels are regulated by autophagy (**Fig. 1a**, **1b**), we next examined if CTR1 is targeted to autophagosomes, the double-membraned autophagic vesicles that deliver cargos to the vacuole for degradation^22^. We used stable transgenic plants expressing GFP- CTR1 from its native promoter in a *ctr1-2* null mutant background (*CTR1p:GFP-CTR1/ctr1-2*). Incubation of GFP-CTR1 seedlings with ConA increased the number of GFP-CTR1-containing vesicles after carbon starvation stress. However, the accumulation of GFP-CTR1 into vesicles was nearly eliminated in the *atg7-2* mutant background, indicating that these vesicles are indeed autophagic vesicles (**Fig. 1c**). To confirm further that the ConA-dependent CTR1- containing vesicles are autophagic vesicles, we analyzed the intracellular colocalization of GFP- CTR1 with mCherry-ATG8a-containing autophagic vesicles in roots of stable transgenic *Arabidopsis* plants (**Fig. 1d**). Manders’ overlap coefficient values revealed that 56.6% of the GFP-CTR1 vesicles colocalized with mCherry-ATG8a vesicles, while 57.2% of mCherry-ATG8a autophagosomes colocalized with vesicles containing GFP-CTR1 (**Fig. 1d**)^38^. These results show that the GFP-CTR1 signal significantly colocalized with mCherry-ATG8a, consistent with the autophagic turnover of CTR1. Bimolecular fluorescence complementation (BiFC) and co-immunoprecipitation analyses further revealed that CTR1 interacts with ATG8a in plants (**Fig. 1e**, **1f**), indicating that CTR1 is either a direct cargo of autophagy or acts as a selective autophagy receptor, mediating the degradation of unknown cargo. Notably, given that CTR1 does not contain a canonical ATG8-interacting motif (AIM) site, this result also suggests that CTR1 interacts with ATG8 through a non-canonical ATG8-binding site^22, 39, 40^. Additionally, CTR1 colocalizes with multiple ATG proteins involved in autophagosome formation, further indicating the functional link between CTR1 and autophagy (**Supplementary** Fig. 1).

### Autophagy positively regulates ethylene responses in *Arabidopsis* by decreasing CTR1 protein levels

The autophagic turnover regulation of CTR1 protein suggests that autophagy modulates ethylene responses in plants. To elucidate the functional relationship between ethylene and autophagy, we examined the ethylene responsiveness of *atg* mutant seedlings under light and dark conditions. Ethylene and its biosynthetic precursor, ACC, have distinct effects on hypocotyl cell expansion in *Arabidopsis* seedlings, stimulating elongation in the light while inhibiting growth in darkness^41, 42^. By examining these morphological responses in *atg* mutants in response to ethylene or ACC, we aimed to determine whether disrupting autophagy perturbs ethylene responses, at least in the context of hypocotyl growth regulation. We selected six *atg* mutants (*atg2*, *atg5*, *atg7*, *atg9*, *atg11*, and *atg18a*), that are defective in autophagy^43^. In the absence of ACC, the hypocotyl lengths of all dark-grown *atg* mutants were significantly longer than those of WT seedlings (**Fig. 2a**). All etiolated *atg* mutants also exhibited a hyposensitive response to a broad range of ACC concentrations compared to the WT, displaying longer hypocotyls than the WT at different ACC concentrations (**Fig. 2b**). The measurement of ethylene biosynthesis in dark-grown *atg* seedlings showed comparable levels to the WT (**Fig. 2c**), suggesting that the altered hypocotyl growth observed in the *atg* mutants is likely due to an altered ethylene response rather than ethylene production. In the absence of ACC treatment, light-grown *atg* mutant seedlings exhibited hypocotyl lengths that were indistinguishable from those of WT seedlings. However, upon exposure to increasing concentrations of ACC, the responsiveness of these mutants to ACC was significantly diminished, leading to considerably shorter hypocotyls compared to WT seedlings (**Fig. 2d, 2e**). No difference in ethylene biosynthesis was observed in light-grown *atg* mutant seedlings compared to the WT (**Fig. 2f**).

**Figure 2.**
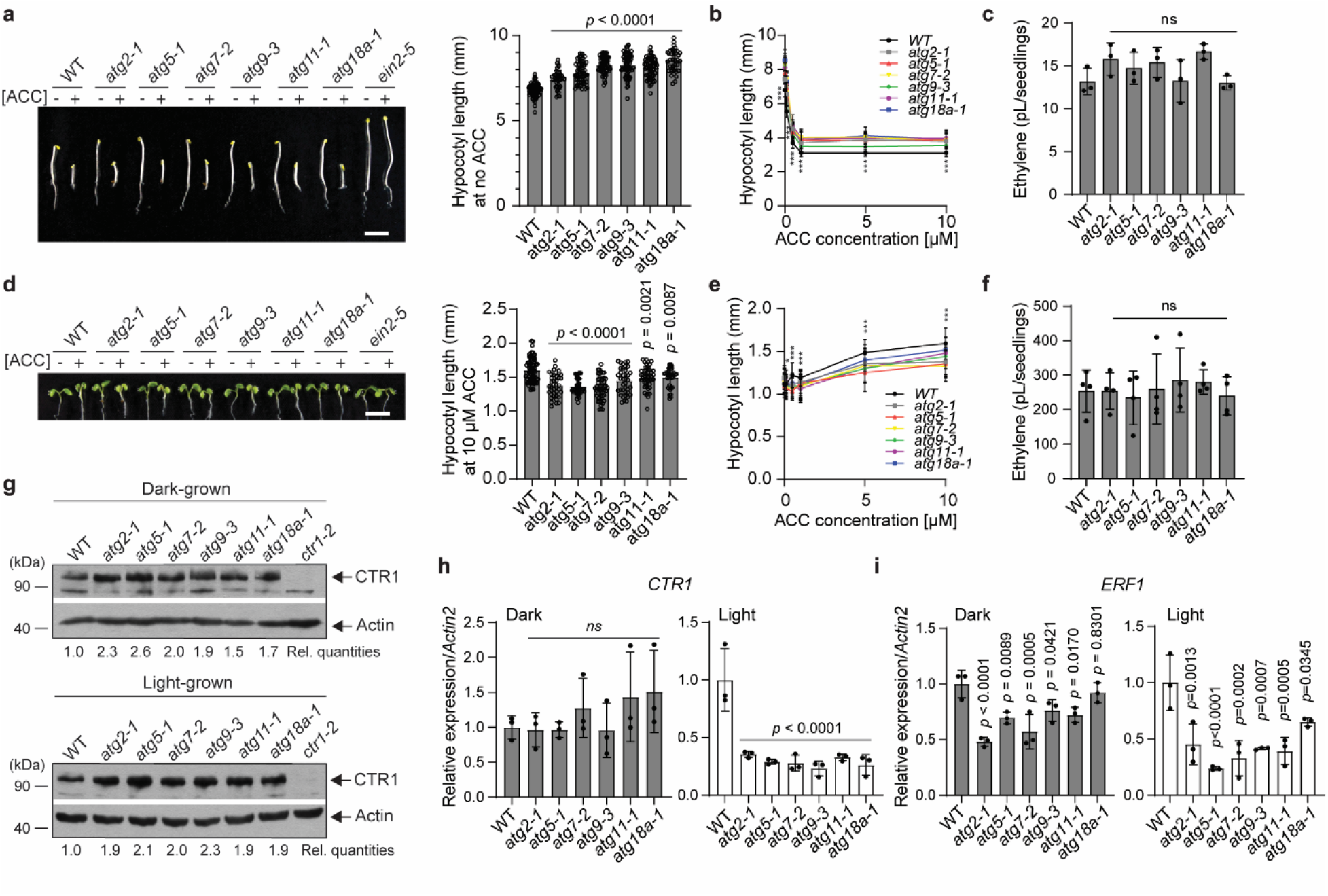
Autophagy positively regulates ethylene responses by modulating CTR1 levels. **a**. *atg* mutants are less sensitive to ACC treatment in darkness. Seedlings were grown on MS medium with or without 10 μM ACC for 3 days in darkness and photographed. The graph presents quantification of the hypocotyl lengths of seedlings grown on MS without ACC. Scale bar, 5 mm. One-way ANOVA with Dunnett’s test was performed to determine statistical significance. Error bars, SD (n ≥ 38 seedlings). **b**. ACC dose-response curves for the hypocotyl length of 3-day-old dark-grown WT and *atg* mutant seedlings. Error bars, SD (n ≥ 38 seedlings for each ACC concentration). Two-tailed Student’s t-test. **c**. Etiolated atg mutant seedlings produced comparable levels of ethylene to the WT seedlings. Seedlings were grown on MS medium for 3 days in darkness, and the accumulated ethylene was measured using gas chromatography. Error bars, SD (n = 3 vials and n ≥ 216 seedlings in each vial). One-way ANOVA with Dunnett’s test was performed to determine statistical significance**. d**. *atg* mutants are less sensitive to ACC treatment in the light. Seedlings were grown on MS medium with or without 10 μM ACC for 5 days in the light and photographed. The graph presents the quantification of the hypocotyl lengths of seedlings grown on MS with 10 μM ACC. Scale bar, 5 mm. One-way ANOVA with Dunnett’s test was performed to determine the statistical significance. Error bars, SD (n ≥ 32 seedlings). **e**. ACC dose-response curves for the hypocotyl length of 5-day-old light-grown WT and *atg* mutant seedlings. Error bars, SD (n ≥ 34 seedlings for each ACC concentration). Two-tailed Student’s t-test. **f**. Light-grown *atg* mutant seedlings produced comparable levels of ethylene to the WT seedlings. Seedlings were grown on MS medium for 5 days in the light, and the accumulated ethylene was measured using gas chromatography. Error bars, SD (n = 4 vials and n ≥ 17 seedlings in each vial). One-way ANOVA with Dunnett’s test was performed to determine the statistical significance. **g**. *atg* mutant expressed higher levels of CTR1 protein compared to WT seedlings. Total protein extracts of 3-day-old etiolated and 5-day-old light-grown seedlings were immunoblotted using an anti-CTR1 antibody. An anti-actin antibody was used as a loading control. Rel. quantities represent the ratio of the intensity of the CTR1 band to actin band signals, and these values are expressed relative to the intensity of CTR1/actin in the WT, which was set to 1. **h, i**. Quantitative gene expression analysis of *CTR1* and *ERF1* in WT and *atg* mutant seedlings. Expression was normalized to an *Actin2* control and is presented relative to the WT control. One-way ANOVA with Dunnett’s test was performed to determine statistical significance. Error bars, SD (n = 3 biological replicates).

Given that autophagy modulates the turnover of the CTR1 protein, the reduced ethylene sensitivity observed in *atg* mutants could partly stem from increased endogenous CTR1 protein levels, which results in an attenuation of ethylene responses. In support of this hypothesis, immunoblot analysis revealed significantly higher CTR1 protein accumulation in the *atg* mutant seedlings than in the WT under both light and dark growth conditions (**Fig. 2g**). However, transcript levels of *CTR1* in the *atg* mutants exhibited either similar or reduced expression relative to those in WT seedlings under dark and light conditions, respectively (**Fig. 2h**). This suggests that the elevated CTR1 protein levels in *atg* mutants result from post-transcriptional effects due to impaired autophagic flux in these mutants. Supporting the enhanced CTR1 protein levels, both dark- and light-grown *atg* mutant seedlings expressed significantly lower levels of the ethylene-inducible *ERF1* than WT seedlings (**Fig. 2i**)^44^.

Ethylene and autophagy have opposing effects on CTR1 turnover (**Fig. 1**)^6, 20^. Ethylene and its precursor ACC increase CTR1 levels, whereas autophagy promotes its protein turnover. To investigate whether autophagy inhibits the ethylene-induced stabilization of CTR1, thereby destabilizing CTR1, we examined the steady-state levels of CTR1 in response to increasing concentrations of ACC in *atg5* and *atg7* mutants, which are representative *atg* mutants with severe autophagy defects. Immunoblot analysis revealed that ACC gradually increases CTR1 levels in a concentration-dependent manner in WT seedlings, corroborating previous reports (**Supplementary** Fig. 2a)^6, 20^. The *atg5* and *atg7* mutants also exhibited a similar pattern of CTR1 accumulation in response to ACC, despite their significantly higher basal CTR1 levels compared to WT seedlings. This suggests that autophagy does not destabilize CTR1 by interfering with ethylene-induced CTR1 stabilization. Instead, an alternative pathway, likely the 26S proteasome-ubiquitin system, may regulate CTR1 turnover. Supporting this notion, treatment with Bortezomib (Btz), a proteasome inhibitor, prevented the degradation of endogenous CTR1 following cycloheximide treatment (**Supplementary** Fig. 2b), demonstrating that CTR1 protein turnover is also regulated by the 26S proteasome pathway. Collectively, these results demonstrate that autophagic flux positively regulates ethylene responses by degrading its key negative regulator, CTR1, thus potentiating ethylene signaling.

### Ethylene signaling mutants exhibit altered carbon starvation-induced autophagic responses

Autophagy-deficient mutants exhibit reduced ethylene responses in both light- and dark-grown *Arabidopsis* seedlings (**Fig. 2**), suggesting the positive regulatory role of autophagy in the ethylene signaling-mediated developmental process. To further elucidate the functional link between autophagy and ethylene, we investigated whether the crosstalk between the two pathways is reciprocal. To this end, we assessed the autophagic responses of constitutive ethylene response and ethylene-insensitive mutants under carbon starvation. The constitutive ethylene response mutant *ctr1-2* showed a hypersensitive response to carbon starvation, exhibiting severe chlorosis compared to WT and failing to recover in the light (**Fig. 3a**). The introduction of a *CTR1* genomic transgene with the *CTR1* promoter driving a *GFP-CTR1* coding fusion complemented the carbon starvation hypersensitivity phenotype of *ctr1-2*, restoring autophagic phenotypes to WT levels. The *ctr1-2* mutant displayed significantly reduced total protein levels and levels of the large subunit of Rubisco (RbcL) seven days after starvation.

**Figure 3.**
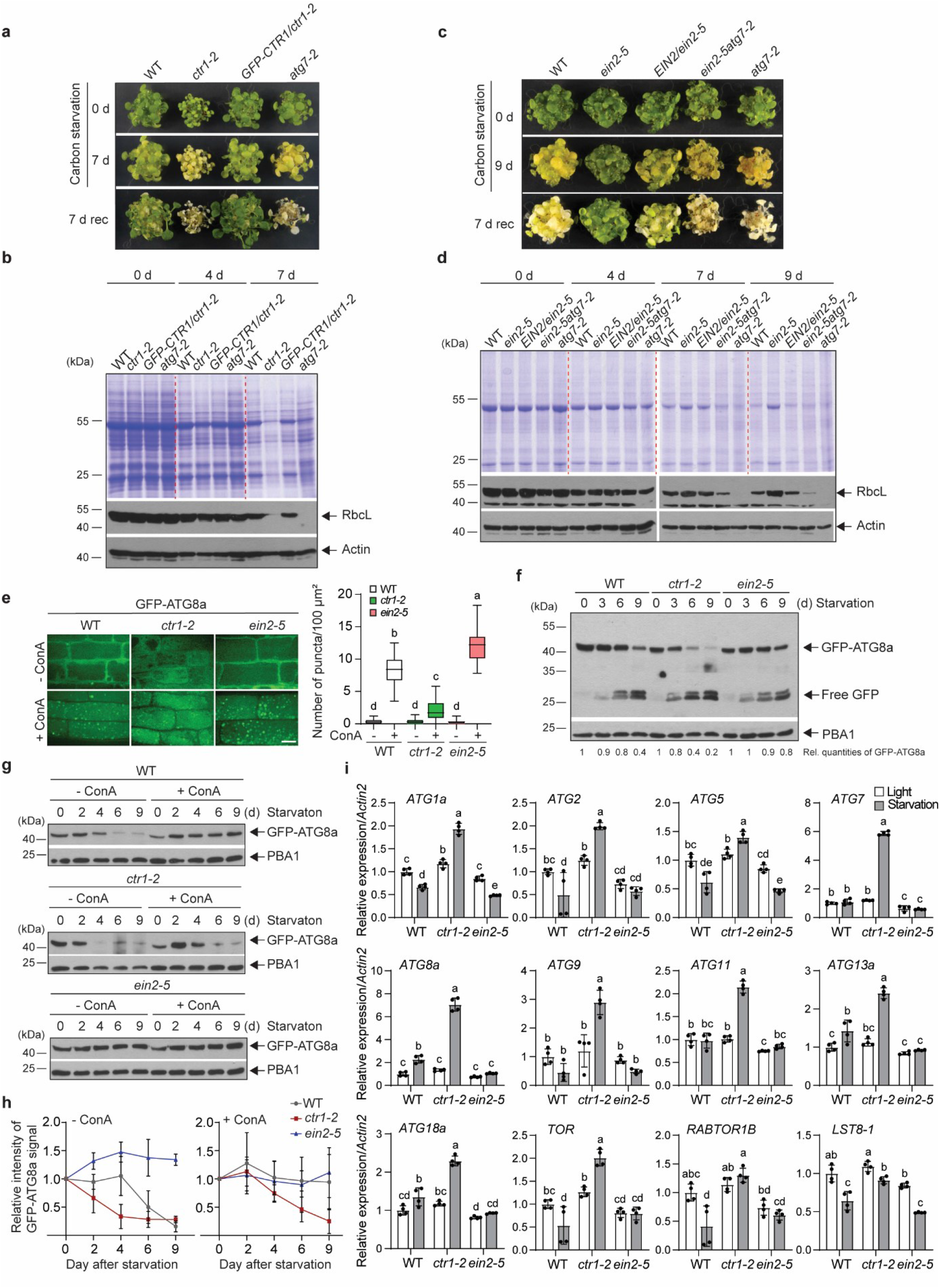
The ethylene signaling pathway negatively regulates autophagic responses under carbon starvation. **a**, Carbon starvation-induced senescence phenotypes of WT, *ctr1-2*, *CTR1p:GFP-CTR1/ctr1-2*, and *atg7-2* mutant seedlings. Seven-day-old light-grown seedlings were grown on MS medium without sucrose and transferred to darkness for 7 days, followed by recovery in the light for 7 days (7-rec). **b**, Total cellular proteins from seedlings during carbon starvation were resolved in SDS-PAGE, stained with Coomassie brilliant blue (CBB), or immunoblotted using antibodies against RbcL and actin (loading control). **c**, Carbon starvation- induced senescence phenotypes of WT, *ein2-5*, *EIN2p:EIN2/ein2-5*, *ein2-5atg7-2*, and *atg7-2* mutant seedlings. **d**. Total cellular proteins from seedlings during carbon starvation were resolved in SDS-PAGE, stained with CBB, or immunoblotted using antibodies against RbcL and actin (loading control). **e**. Autophagosome formation is altered in ethylene signaling mutants. Five-day-old light-grown seedlings expressing GFP-ATG8a in WT, *ctr1-2*, or *ein2-5* were incubated in a sucrose-free medium with and without ConA under light or dark conditions for 16 hours. Root cells were examined for autophagic vesicles by confocal microscopy. Scale bars, 10 μm. One-way ANOVA with Tukey’s post-hoc test was performed, and different letters denote statistically significant differences between groups, *p* ≤ 0.0007. (n ≥ 20 cells from five to six seedlings for each treatment). **f**. Time course of free GFP release from GFP-ATG8a during carbon starvation. WT, *ctr1-2* and *ein2-5* seedlings expressing GFP-ATG8a were carbon starved for the indicated times. Total protein extracts were immunoblotted with anti-GFP antibodies. Arrows indicate the GFP-ATG8a fusions and the free GFP released upon degradation. Relative quantities represent the ratio of the GFP-ATG8a band intensity to the PBA1 band intensity. For each genotype, these ratios are normalized to the GFP-ATG8a/PBA1 ratio in the corresponding no-carbon starvation control, which is set to 1. **g**. The steady-state levels of GFP-ATG8a in WT, *ctr1-2*, and *ein2-5* mutants. Total protein extracts of seedlings after the indicated starvation were immunoblotted using anti-GFP and anti-PBA1 antibodies. **h**. Quantification of GFP-ATG8a bands during starvation with and without ConA. Relative quantities of GFP-ATG8a bands are represented as the ratio of the GFP-ATG8a band intensity to PBA1 band intensity. For each genotype, these ratios are normalized to the GFP- ATG8s/PBA1 ratio in the corresponding no starvation control, which is set to 1. Error bars, SD (3 biological replicates). **i**, Quantitative gene expression analysis of *ATGs*, *TOR*, *Raptor1b*, and *LST8-1* genes in WT, *ctr1-2*, and *ein2-5* mutant seedlings. Expression was normalized to an *Actin*2 control and is presented relative to the WT control. One-way ANOVA with Tukey’s post- hoc test was performed, and the different letters denote statistically significant differences between groups. Error bars, SD (n = 4 biological replicates).

These defects were fully alleviated in the complementation lines harboring the *CTR1* transgene, with total protein and RbcL levels restored to WT levels (**Fig. 3b**). In contrast, the ethylene-insensitive *ein2-5* mutant displayed significantly delayed leaf chlorosis, which was sustained even after nine days in darkness (**Fig. 3c**). The *ein2-5* mutant continued to express higher levels of total cellular proteins and RbcL protein compared to WT under carbon starvation.

Complementation of *ein2-5* by expressing a genomic *EIN2* transgene rescued the carbon starvation insensitivity of *ein2-5* to WT levels (**Fig. 3c, 3d**), confirming that EIN2 is responsible for the carbon starvation-insensitive phenotypes of *ein2-5*. Furthermore, introducing an *atg7* mutation into *ein2-5* (*ein2-5atg7-2* double mutant) inhibited the carbon starvation insensitivity of *ein2-5*, positioning autophagy downstream of ethylene signaling (**Fig. 3c**, **d**). Similar to *ein2-5*, the ethylene-insensitive mutants *ein3eil1* and *etr1-1* exhibited hyposensitive carbon starvation responses and maintained higher levels of total cellular proteins and RbcL compared to WT seedlings under carbon starvation conditions (**Supplementary** Fig. 3).

### Ethylene signaling negatively regulates autophagy likely through destabilizing autophagosomes and modulating transcription

The dynamic engulfment and delivery of cellular components to the vacuoles via autophagosomes is a key process in autophagy^22^. To investigate the influence of ethylene signaling on autophagosome formation in response to carbon starvation, we examined this process in *ein2-5* and *ctr1-2* mutants using GFP-ATG8a reporter lines expressing GFP-ATG8a in the WT, *ctr1-2*, or *ein2-5* mutant backgrounds. We selected *ein2-5*, *ctr1-2*, and WT lines with similar GFP-ATG8a protein levels for the carbon starvation experiments (**Supplementary** Fig. 4). Carbon starvation treatment of *35S:GFP-ATG8a/WT* seedlings in the presence of ConA led to an elevated number of GFP-ATG8a-labeled autophagosomes compared to the ConA untreated control. This increase in autophagosome formation was even more pronounced in *35S:GFP-ATG8a/ein2-5* seedlings under the same starvation conditions compared to the WT (**Fig. 3e**). Conversely, *35S:GFP-ATG8a/ctr1-2* mutants exhibited only a marginal rise in the quantity of GFP-ATG8a autophagosomes when exposed to carbon starvation. (**Fig. 3e**).

Additionally, treatment with benzothiadiazole (BTH), a salicylic acid analog that induces autophagy, further increased autophagic flux in *35S:GFP-ATG8a/WT* and *35S:GFP- ATG8a/ein2-5* seedlings after carbon starvation in the presence of ConA^45, 46^. However, this increase was not observed in *35S:GFP-ATG8a/ctr1-2* mutants (**Supplementary** Fig. 5). Next, we investigated whether ethylene signaling influences autophagic flux under carbon starvation conditions. Intriguingly, despite the contrasting effects on the number of GFP-ATG8a-labeled autophagosomes in the *ctr1-2* and *ein2-5* mutants compared to WT (**Fig. 3e**), GFP cleavage assays revealed a striking similarity in the release kinetics and amounts of free GFP among the mutants and WT after starvation (**Fig. 3f**). However, the steady-state levels of intact GFP- ATG8a significantly differed between *ctr1-2* and *ein2-5* compared to WT. In *ctr1-2*, GFP-ATG8a degraded faster upon starvation, while no apparent decrease in GFP-ATG8a levels was observed in *ein2-5* (**Fig. 3f**). These results suggest that ethylene signaling does not control autophagosome degradation in the vacuole. Rather, the *ctr1-2* mutation likely either accelerates ATG8 protein degradation or impedes autophagosome formation before vacuolar fusion, leading to fewer autophagosomes. To test these possibilities, we subjected *35S:GFP-ATG8a/WT*, *35S:GFP-ATG8a/ctr1-2,* and *35S:GFP-ATG8a/ein2-5* seedlings with or without ConA treatment during carbon starvation (**Fig. 3g**). ConA inhibits vacuolar (H+)-ATPases, thus preventing the degradation of autophagic cargos within vacuoles. In WT seedlings, ConA treatment maintained constant levels of GFP-ATG8a during starvation compared to untreated controls. However, in *ctr1-2* mutants, GFP-ATG8a levels initially stabilized briefly but then rapidly degraded, similar to untreated samples. In contrast, *ein2-5* mutants exhibited constant GFP-ATG8a levels during starvation, unaffected by ConA treatment (**Fig.3g**, **3h**). To further investigate whether CTR1 directly regulates ATG8a stability, we examined the steady-state levels of endogenous ATG8a in *ctr1-2* mutants and transgenic lines overexpressing GFP-CTR1 (CTR1ox) (**Supplementary** Fig. 6). Immunoblot analysis showed no significant differences in ATG8a levels among WT, *ctr1-2*, and CTR1ox lines with and without autophagy-inducer BTH treatment. This implies that CTR1 does not significantly impact overall ATG8a protein abundance. Instead, the role of CTR1 appears to be more specific to autophagosome stabilization or turnover. Collectively, these findings suggest that ethylene signaling inhibits autophagy by destabilizing autophagosomes before vacuolar fusion, while CTR1 counteracts this process.

To elucidate the molecular mechanism by which ethylene signaling negatively regulates autophagy at the transcriptional levels, we examined the transcriptional regulation of genes involved in autophagy and the TOR signaling pathway, which is a well-known negative regulator of autophagy^47, 48^. Under normal conditions, quantitative RT-PCR analysis showed similar transcript levels of various *ATG* genes in WT, *ctr1-2*, and *ein2-5* mutants. However, upon carbon starvation, *ctr1-2* mutants exhibited significantly higher expression of most *ATG* genes compared to WT and *ein2-5,* while WT and *ein2-5* showed reduced or similar transcript levels of several *ATG* genes (**Fig. 3i**). Intriguingly, the transcript levels of *TOR*, *RAPTOR1B*, and *LST8-1*, which encode components of the TOR signaling pathway, were also significantly higher in *ctr1-2* mutants after carbon starvation, with particularly notable increase in *TOR* and *RAPTOR1B* (**Fig. 3i**). The elevated expression of most *ATG* genes in *ctr1-2* mutants likely represents a compensatory response to impaired autophagosome formation in an attempt to upregulate autophagy-related pathways to counteract the reduced autophagic activity. Concurrently, the upregulated expression of TOR signaling components supports the observed reduction in autophagic activity in *ctr1-2* mutants. These results demonstrate that ethylene signaling also regulates autophagy in part through transcriptional regulation targeting key autophagy and TOR components, revealing an interplay between ethylene signaling, autophagy, and TOR pathway regulation.

### Serine 924 of EIN2 plays a key role in regulating autophagic responses to carbon starvation

The constitutive ethylene response mutant *ctr1-2* showed hypersensitive responses to carbon starvation. Thus, we further sought to elucidate the autophagic responses of constitutive nuclear localization of EIN2 and the role of Serine 645 (S654) and Serine 924 (S924), both of which lead to the cleavage of EIN2 and subsequent activation of ethylene signaling^12, 13, 49^. To investigate this, we examined carbon starvation phenotypes of transgenic *Arabidopsis* seedlings expressing WT full-length EIN2 (EIN2^WT^) or an *EIN2* genomic fragment with S654A (EIN2^S645A^), S924A (EIN2^S924A^), or both substitutions (EIN2^AA^) under its native promoter in the *ein2-5* background (**Fig. 4a**)^12^. The *EIN2p-EIN2^WT^* transgene alleviated the delayed leaf senescence of the *ein2-5*. In contrast, both *EIN2p-EIN2^AA^*and *EIN2-EIN2^S924A^* transgenes were unable to rescue the carbon starvation phenotypes of the *ein2-5*, instead conferring hypersensitive carbon starvation responses similar to those observed in *ctr1-2* mutants. Notably, the *EIN2p-EIN2^S645A^* transgene fully complemented the phenotypes observed in the *ein2-5* mutant. Under carbon starvation, *EIN2p-EIN2^S645A^*plants maintained total protein and RbcL accumulation levels comparable to those of WT. In contrast, carbon-starved *EIN2p-EIN2^S924A^*plants exhibited severe depletion of protein and RbcL levels, akin to the patterns seen in the *atg7-2* mutant (**Fig. 4b**). These observations suggest that CTR1-mediated phosphorylation of EIN2 at S924, but not S645, plays a pivotal role in suppressing autophagy during carbon starvation conditions.

**Figure 4.**
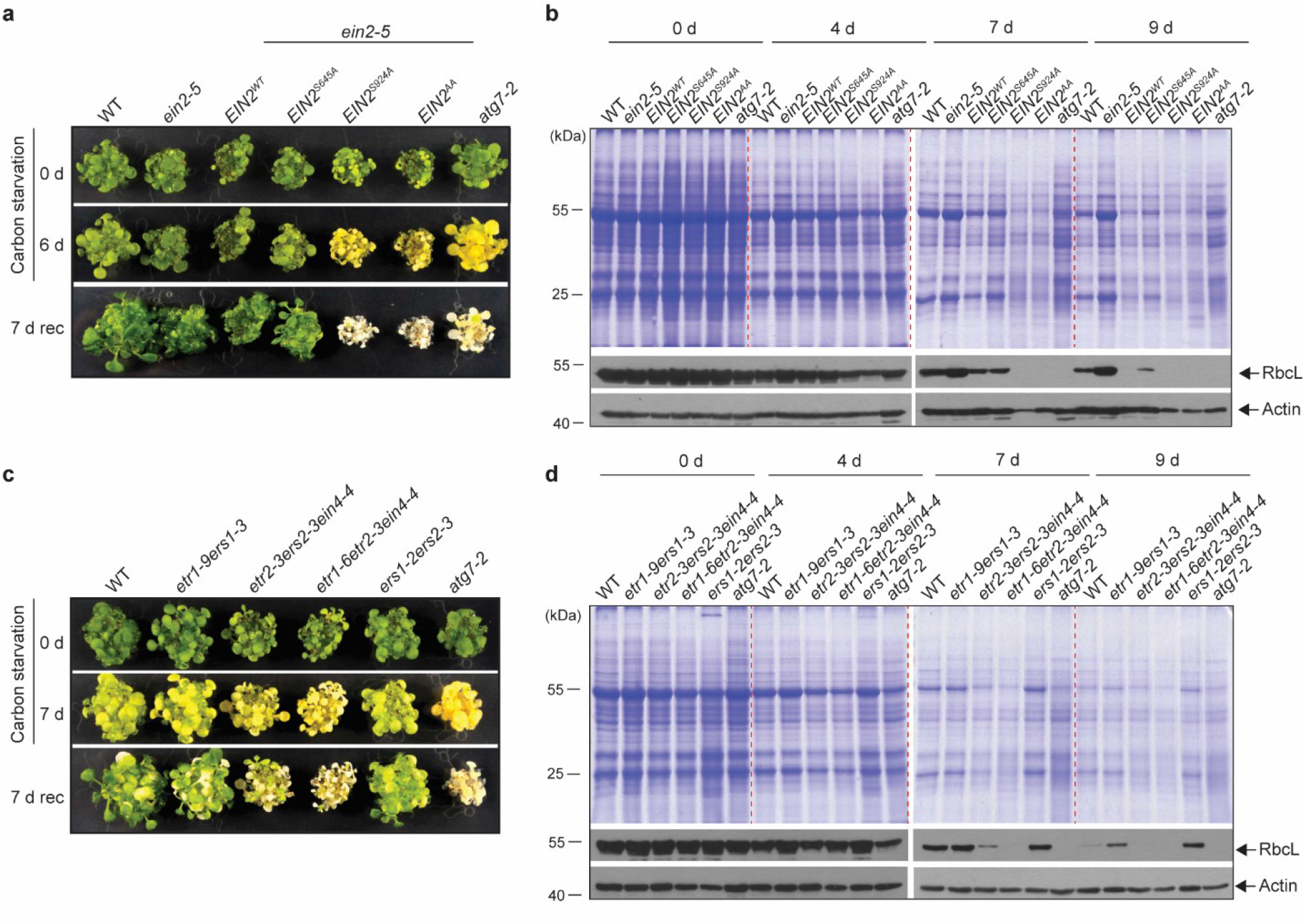
CTR1-mediated EIN2 phosphorylation and ethylene receptors with a receiver domain play a role in autophagic responses during carbon starvation. **a**. Carbon starvation-induced senescence phenotypes of WT, *ein2-5*, and *ein2-5* complementation lines expressing *EIN2^WT^*, *EIN2^S645A^*, *EIN2^S924A^*, or *EIN2^AA^* under its native promoter. Seven-day-old light-grown seedlings were grown on MS medium without sucrose and transferred to darkness for 7 days, followed by recovery in the light for 7 days (7-rec). **b**. Total cellular proteins from seedlings during carbon starvation were resolved in SDS-PAGE, stained with CBB, or immunoblotted using antibodies against RbcL and actin (loading control). **c**. Carbon starvation- induced senescence phenotypes of WT and ethylene receptor mutants with or without a receiver domain or kinase activity. **d**. Total cellular proteins from seedlings during carbon starvation were resolved in SDS-PAGE, stained with CBB, or immunoblotted using antibodies against RbcL and actin (loading control).

### Ethylene receptors with a receiver domain play a major role in regulating autophagy independently of kinase activity

CTR1 localizes to the ER as part of an ethylene receptor signaling complex through its direct interaction with the C-terminal domain of the receptors, which consists of a kinase domain with or without a receiver domain^20, 50^. The ER-localized ethylene receptors consist of five members: Ethylene Response 1 (ETR1), Ethylene Response Sensor 1 (ERS1), ETR2, ERS2, and EIN4. These receptors are classified into subfamilies I (ETR1 and ERS1) and II (ETR2, EIN4, and ERS2) based on the number of transmembrane domains and the homology of the histidine kinase domain ^8, 51–55^. Subfamily I members primarily exhibit histidine kinase activity, whereas subfamily II members may possess serine/threonine kinase activity. Of the five receptors, ETR1, ETR2, and EIN4 have a receiver domain as part of their C-terminus, while ERS1 and ERS2 lack this domain^53, 56, 57^. To investigate the role of the receiver domain and receptor histidine kinase activity in autophagy during carbon starvation, we examined the autophagic phenotypes of various ethylene receptor mutant combinations (**Fig. 4c**). The *etr1-9ers1-3* mutant (lacking receptors with histidine kinase activity) and the *ers1-2ers2-3* mutant (lacking receptors without a receiver domain) exhibited a WT-like chlorosis phenotype and reduced total protein abundance and RbcL levels in response to carbon starvation (**Fig. 4d**)^58, 59^. In contrast, the *etr1-6etr2-3ein4- 4* mutant (lacking receptors with a receiver domain) exhibited hypersensitivity to starvation similar to *ctr1-2* and *atg7-2* mutants^60^. The *etr2-3ers2-3ein4-4* triple mutant (lacking subfamily II receptors) also showed hypersensitivity, but with a lesser reduction in RbcL and total protein compared to the *etr1-6etr2-3ein4-4* (**Fig. 4d**). These results suggest that receptors without a receiver domain are non-essential, while receptors with a receiver domain play a primary role in autophagy, regardless of kinase activity. Consistent with this, *etr1-6* and *etr1-7* loss-of-function mutants also displayed hypersensitive responses to carbon starvation (**Supplementary** Fig. 7)^61, 62^. Together, these results indicate that ethylene receptors with a receiver domain are essential for the autophagic response to carbon starvation, presumably via interaction with CTR1 through the receiver domain, while receptor kinase activity is nonessential^63^.

## Discussion

CTR1 has long been recognized as a critical regulator of the ethylene signaling pathway. However, how its protein level is controlled has remained enigmatic. In this study, we demonstrate that autophagy degrades CTR1 proteins in response to carbon-deficient stress, identifying autophagy as a key regulator of ethylene responses by controlling CTR1 levels under stress conditions. We also reveal reciprocal crosstalk between ethylene signaling and autophagy mediated by autophagic CTR1 destabilization, wherein autophagy stimulates ethylene responses, while ethylene signaling inhibits autophagy. Our findings unveil that autophagy-mediated CTR1 degradation is a key mechanism linking and coordinating these two signaling networks that govern plant growth and stress responses.

While the biochemical and physiological stress responses of ethylene and autophagy pathways are well known, how plants balance these signaling pathways to control the magnitude and duration of stress responses is unclear. Preventing overshoot stress responses is crucial for avoiding plant damage and death while maintaining the capacity to resume growth after stress relief. Our finding that ethylene and autophagy reciprocally regulate each other through modulation of CTR1 levels provides insight into these balancing stress responses and proposes a model for the crosstalk between the two pathways (**Fig. 5**). When ethylene signaling becomes overactive, it suppresses autophagy, which is attenuated by the TOR kinase cascade under normal growth conditions. This suppression leads to the accumulation of CTR1, consequently alleviating further intensification of ethylene responses. Conversely, hyperactivated autophagy promotes the degradation of CTR1, releasing ethylene signaling from inhibition to restrain autophagic activity. Thus, CTR1 enables each pathway to shut off the other upon exceeding moderate thresholds while permitting balanced crosstalk to sustain optimal stress responses. Collectively, by leveraging CTR1 to modulate both pathways, ethylene signaling and autophagy may be able to activate protective stress responses while still preventing excessive responses that compromise the long-term viability of plants. This elegant homeostatic feedback loop modulates transient and defensive stress responses, followed by a coordinated recovery of growth homeostasis, unveiling molecular strategies to enhance environmental stress resilience in plants.

**Figure. 5.**
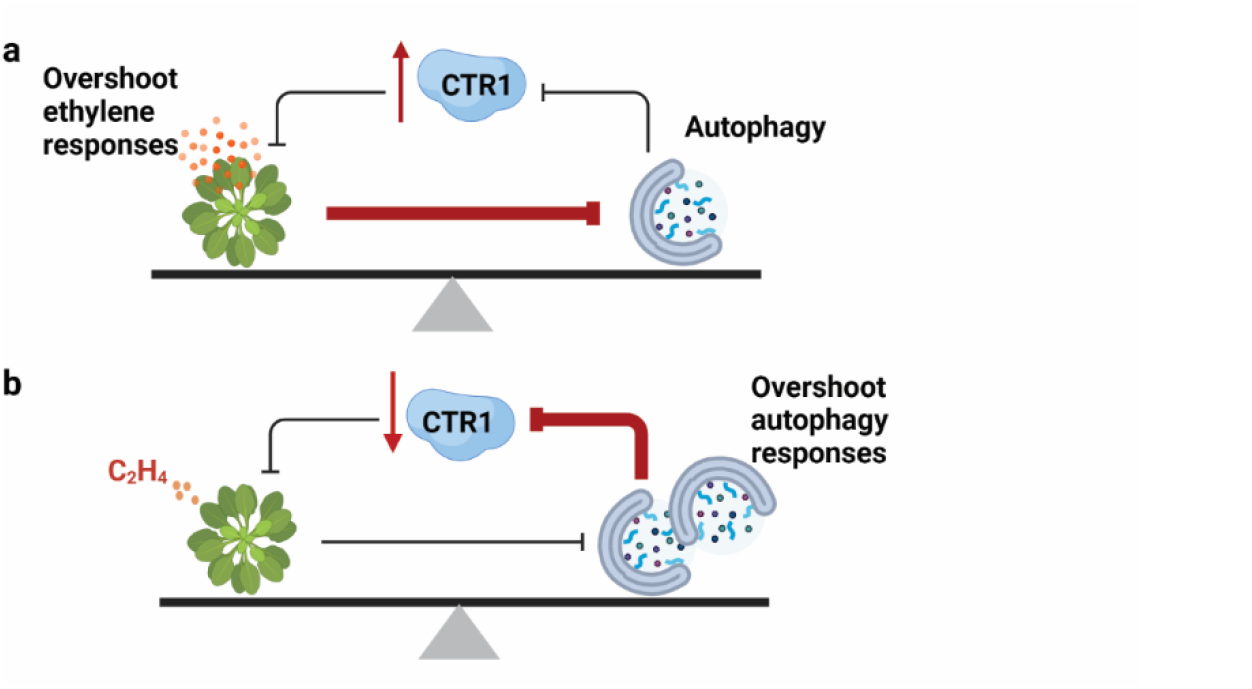
Model for the reciprocal regulation between ethylene signaling and autophagy. The survival of plants under stress conditions hinges on maintaining a delicate balance between mounting stress responses and preserving their recovery potential. Plants achieve this balance through the reciprocal regulation of ethylene signaling and autophagy with CTR1 acting as a crucial crosstalk node to prevent an overshoot of stress responses. By modulating CTR1 levels, plants can mitigate excessive activation of either pathway, thereby preventing an overshoot of stress responses while retaining the ability to recover once the stress has subsided. **a**. When ethylene signaling becomes overactivated, it suppresses autophagy, subsequently inhibiting the autophagy-mediated turnover of CTR1. The resulting accumulation of CTR1 then dampens the ethylene response, forming a negative feedback loop that mitigates the overshoot of ethylene signaling. **b**. Conversely, excessively activated autophagy further stimulates CTR1 turnover, leading to an inhibition of the CTR1-mediated constraint on ethylene signaling. The enhanced ethylene response then attenuates autophagy activity, ultimately mitigating the overactivation of autophagy. Through these interconnected negative feedback inhibition pathways, ethylene signaling and autophagy counterbalance each other when either process becomes dominant. This intricate interplay defends the plants against runaway activity from either a critical stress response or an excessive recovery program, ensuring their survival. The thickness of the lines represents the strength of inhibition, with thicker lines indicating stronger inhibition.

The CTR1-mediated phosphorylation of EIN2 regulates the proteolytic cleavage of EIN2, transducing ethylene signals from the ER to the nucleus^12, 13, 49^. Given the hypersensitive and insensitive autophagic responses of *ctr1-2* and *ein2-5* mutants, respectively, to carbon starvation, activated ethylene signaling through proteolytic cleavage of EIN2 regulated by two phosphorylation sites, S645 and S924, appears essential for suppressing autophagy. However, unlike *EIN2^S924A^* plants, which showed hypersensitive autophagy as expected, the S645A mutation did not confer a hypersensitive autophagic response to carbon starvation. This result somewhat echoes the substantially weaker ethylene responses of *EIN2^S645A^* plants compared to *EIN2^S924A^* plants in inhibiting leaf cell expansion of light-grown and etiolated seedlings^12^. The differential autophagy impacts of S645A and S924A mutations likely stem from multifaceted regulatory mechanisms. The absence of S645A effects on autophagy raises questions about potential redundancy from the modification of additional residues obscuring S645A effects. Notably, the TOR kinase phosphorylates EIN2 at S657, blocking EIN2 nuclear translocation and ethylene signaling activation^64^. Given the close proximity of S645 to S657 and the interplay between autophagy and TOR signaling, TOR-mediated regulation of EIN2 may compensate for the S645A mutation. Specifically, TOR-induced phosphorylation of EIN2 at S657 could potentially mask the effects of the S645A mutation, inhibiting EIN2 nuclear translocation. This compensatory mechanism may account for the higher expression of transcripts encoding TOR components observed in *ctr1-2* mutants (**Fig. 3i**), representing a negative feedback loop to inhibit ethylene signaling when the EIN2 function is highly activated. Moreover, S645 and S924 may transduce distinct downstream signals, with S924 directly influencing autophagy machinery and S645 affecting peripheral pathways. Finally, localization or stability changes from the mutations could alter EIN2 activity. Further elucidating site-specific signaling outcomes and dissecting this intricate phosphorylation-based control of EIN2 and autophagy will be essential to explain the divergence between S645A and S924A mutations.

CTR1 interacts with ATG8 despite lacking a canonical ATG8-interacting motif (AIM) (**Fig. 1e** and **f**). This suggests that non-canonical binding sites may facilitate CTR1-ATG8 interaction. Intriguingly, CTR1 contains a motif (EESYQLQLALALRLSSE) with high sequence homology to the ubiquitin-interacting motif (UIM), a novel class of alternative ATG8 interaction sites that bind the UIM-docking site (UDS) present in ATG8 proteins^39^. The identification of the conserved UIM-UDS interface expands the range of selectively targeted autophagy cargoes that can be recruited independently of AIM sites^39^. The putative UIM-like sequence in CTR1 could facilitate its previously unknown interaction with ATG8 through this alternative UIM-UDS binding mode, providing a mechanism for the AIM-independent autophagic turnover of CTR1. Further biochemical and genetic characterization are needed to confirm whether this putative UIM motif directly mediates CTR1-ATG8a binding via the predicted UDS interface.

The seemingly contradictory results between GFP-ATG8a cleavage assays and the hypersensitive carbon starvation response in the *ctr1-2* mutant (**Fig. 3**) suggest inefficient autophagy that leads to inadequate recycling of cellular components in the *ctr1-2*. GFP cleavage assays revealed similar release kinetics of free GFP across WT, *ctr1-2*, and *ein2-5* backgrounds, indicating that autophagosome degradation proceeds similarly once initiated in all genotypes. However, while ConA treatment stabilized GFP-ATG8a in WT seedlings, it only led to a transient increase in GFP-ATG8a levels in *ctr1-2* seedlings, followed by rapid degradation under starvation. In contrast, *ein2-5* mutants maintained consistent GFP-ATG8a levels regardless of ConA treatment (**Fig. 3g**, **3h**). These results suggest that CTR1 may stabilize ATG8a proteins, and ethylene signaling counteracts this process. However, endogenous ATG8a levels did not significantly differ among WT, CTR1ox, and *ctr1-2* mutants (**Supplementary** Fig. 6), indicating that the role of CTR1 is specific to autophagosome stabilization rather than affecting ATG8a protein abundance. CTR1 colocalizes with multiple ATG proteins in addition to ATG8a (**Supplementary** Fig. 1), implying that it may participate in early autophagosome formation by potentially phosphorylating these proteins, similar to how mammalian mTORC1 kinase influences autophagy through phosphorylation of ATG proteins such as ATG1 and ATG13^65, 66^. These findings propose a model where CTR1 stabilizes autophagosomes, preventing their premature degradation, while ethylene signaling promotes autophagosome destabilization, thereby negatively regulating autophagy. This complex interplay between stabilization and destabilization processes likely determines the overall efficiency of autophagy under carbon starvation conditions.

The differential gene expression regulation of *ATGs* and TOR signaling components in *ctr1-2* and *ein2-5* mutants compared to WT suggests the presence of compensatory mechanisms. Under carbon starvation conditions, *ctr1-2* mutants exhibited increased transcript levels of TOR components, including *TOR* and *RAPTOR 1B*, indicating enhanced TOR signaling activity. Simultaneously, *ctr1-2* expressed higher levels of *ATG* genes compared to WT and *ein2-5* mutants during starvation (**Fig. 3i**). This upregulation of *ATG* gene expression suggests a positive feedback mechanism to enhance autophagy-related processes, possibly in reaction to stress conditions. However, the hypersensitivity of *ctr1-2* mutants to carbon starvation implies that increased *ATG* gene expression alone may not confer effective stress tolerance. We hypothesize that elevated TOR signaling in *ctr1-2* likely inhibits autophagy initiation by phosphorylating and inactivating key autophagy regulators such as Atg13 and ULK1, despite increased expression of *ATG* genes ^65, 67^. Together with the potential role of ethylene signaling in destabilizing autophagosomes, the enhanced TOR signaling in *ctr1-2* could partly explain its hypersensitivity to carbon starvation and decreased autophagosome numbers, despite showing similar levels of autophagic flux markers. Furthermore, the role of TOR signaling in coordinating nutrient responses and growth may disrupt cellular metabolic balance, potentially exacerbating the sensitivity of *ctr1-2* mutants to nutrient deprivation. Further research is needed to understand the specific mechanisms through which ethylene and TOR signaling modulate autophagy and their implications for cellular adaptation to stress.

## MATERIALS AND METHODS

### Plant materials and growth conditions

*Arabidopsis thaliana* Col-0 was used as the WT reference throughout the study. All plants were grown in long-day conditions at 22 °C ± 2 °C or in vitro on half-strength Murashige and Skoog (MS) basal medium supplemented with 0.8 % plant agar (pH 5.7) with or without sucrose in a continuous light chamber at 22 °C ± 2 °C. All transgenic plants were homozygous and were identified by the segregation of specific antibiotic resistance, followed by the confirmation of protein expression via immunoblot analyses. *Arabidopsis* mutants and transgenic plants used in this study include *ctr1-2, ein2-5*, *ein2-5atg7-2*, *etr1-1*, *etr1-6*, *etr1-7*, *ein3eil1*, *etr1-9ers1-3*, *etr1- 2ers2-3*, *etr1-6etr2-3ein4-4*, *etr2-3ers2-3ein4-4*, *atg2-1*, *atg5-1*, *atg7-2*, *atg9-3*, *atg11-1*, *atg18a-1*, *EIN2p:EIN2^WT^/ein2-5*, *EIN2p:EIN2^S645A^/ein2-5*, *EIN2p:EIN2^S924A^/ein2-5, EIN2p:EIN2^AA^/ein2-5*, *CTR1p:GFP-CTR1/ctr1-2*, *CTR1p:GFP-CTR1/atg7-2*, *CTR1p:GFP-CTR1;35S:mCherry- ATG8a/ctr1-2*, *35S:GFP-ATG8a/WT*, *35S:GFP-ATG8a/ctr1-2*, *35S:GFP-ATG8a/ein2-5*, *35S:GFP-CTR1/WT*.

### Plasmid construction

All molecular cloning was performed using Infusion cloning (Takara Bio USA) and Gateway (Invitrogen) strategies. The creation of the *CTR1p-GFP-gCTR1* construct was described in Park et al. (2023). To create the *35S:GFP-CTR1* construct, coding sequences of full-length CTR1 were cloned into the pENTR entry vector (Invitrogen) and subsequently moved to the binary *pSITE-2CA* Gateway vector. To generate the *35S:Myc-ATG8a*, *35S:GFP-ATG8a* and *35S:mCherry-ATG8a* constructs, coding sequences of full-length *ATG8a* were cloned into the pENTR vector and transferred to either binary *pEarleygate 203*, *pSITE-2CA,* or a modified *pEarleygate 100* with mCherry, respectively. To generate the constructs for BiFC, the coding sequences of CTR1, ATG8a and CIP8 in the pENTR Gateway entry vector were transferred into the pCL112 or pCL113 BiFC binary vector.

### Hypocotyl measurements

To measure the hypocotyl lengths of seedlings in response to ACC, *Arabidopsis* Col-0 and autophagy mutants were planted on an MS medium containing different concentrations of ACC. After cold treatment for 72 hours at 4 °C in darkness, the plates were placed vertically at 22 °C in either darkness or light for 3 or 5 days, respectively. The seedlings were photographed at the indicated times, and the hypocotyl length was measured using Image J software.

### Confocal microscopy

Imaging of GFP, YFP, RFP, and mCherry fluorescence was performed using either a laser-scanning confocal microscope (Zeiss LSM880 upright) or a spinning-disk confocal microscope.

Samples of *N. benthamiana* leaves were directly mounted on glass slides in water, and epidermal cells were imaged by merging z-stack images acquired at 2 µm intervals. *Arabidopsis* samples were mounted similarly, and cells within the root elongation zone were imaged. For imaging *Arabidopsis* stable transgenic seedlings expressing GFP-ATG8a or co-expressing GFP-CTR1 and mCherry-ATG8a, seedlings were grown in liquid Gamborg (GM) medium under light for 5 days. For ConA treatment, seedlings were washed three times in liquid GM medium without sucrose and treated with 0.5 µM ConA dissolved in DMSO in the dark for 16 hours. Imaging was performed using a spinning-disk confocal system consisting of an Olympus IX-83 microscope equipped with a Yokogawa scanning unit (CSU-X1-A1; Hamamatsu Photonics) and an Andor iXon Ultra 897BV EMCCD camera (Andor Technology). The root elongation zone was imaged with a 60x (1.40 NA) or 100x (1.45 NA) oil objective (UPlanSApo; Olympus), and z-series were collected at 0.5 µm step sizes. For BiFC imaging, *N. benthamiana* leaves were co-infiltrated with *Agrobacterium* strains expressing nYFP-CTR1, bZIP-RFP, and either cYFP-ATG8a or cYFP-CIP8a. For colocalization analysis, leaves were co-infiltrated with strains expressing GFP-CTR1 and mCherry-ATG8s. After a 3-day incubation in constant light, leaf epidermal cells were imaged for YFP, GFP, RFP, and/or mCherry signals using a Zeiss LSM880 upright confocal microscope. For all multi-channel imaging experiments, cells expressing single fluorescent markers were imaged initially to ensure no fluorescence bleed- through occurred between channels.

### Colocalization analysis

Spatial colocalization of GFP and mCherry fluorescent signals from labeled autophagosomes was quantified using Manders’ overlap coefficient analysis implemented in the JaCoP (Just Another Colocalization Plugin) plug-in for ImageJ ^38^. Regions of interest containing fluorescently labeled autophagosomes were cropped from the acquired images. The cropped images were processed with the automatic thresholding function of JaCoP to remove background signals.

Colocalization percentages were then determined by comparing the GFP (green) images with the mCherry (red) images, and vice versa. To account for random colocalization events, a rotation control was performed. One image from each pair was rotated 180 degrees, and the colocalization analysis was repeated on the rotated and original images to quantify the expected level of colocalization due to chance overlap.

### Measurements of ethylene production

Surface-sterilized seeds were placed on 3 mL of half-strength Murashige and Skoog (MS) medium supplemented with 1% sucrose and 0.8% agar in 22 mL gas chromatography (GC) vials. After 3 day-stratification at 4°C in the dark, the vials were capped, and the seedlings were incubated in the dark for an additional three days. The accumulated ethylene was then measured using a Shimadzu GC2010 Plus capillary gas chromatography system equipped with an HS-20 headspace autosampler. To assess ethylene production in light-grown seedlings, stratified seeds were transferred to light conditions and grown for 4 days, followed by capping and incubation in the light for 2 days. Ethylene concentration was calculated as pL per seedling for 2-day incubation periods. All genotypes and treatments were evaluated using three biological replicates with the average and standard deviation (SD) were calculated.

### Co-immunoprecipitation analysis

*N. benthamiana* leaves were infiltrated with Agrobacteria co-transformed with the plasmids of interest and incubated for 3 days in a growth chamber. Total proteins were extracted from the infiltrated leaves and homogenized in a co-immunoprecipitation buffer (25 Tris, pH 7.5, 150 mM NaCl, 5 mM EDTA, pH 8.0, 0.1% Tween-20, 1 mM DTT, 40 µM MG132, 1 mM PMSF, and 0.5x protease inhibitor cocktail) by a 10-min centrifugation at 5,000 RPM. Following the centrifugation, the supernatant was removed, and the proteins were homogenized and extracted from the pellets in a RIPA buffer (50 mM Tris, pH 7.5, 50 mM NaCl, 2 mM EDTA, pH 8.0, 0.1%

SDS, 5 mM DTT, 40 µM MG132, 1 mM PMSF, 1x protease inhibitor cocktail) by a 10-min centrifugation at 7,000 RPM. After quantifying the protein concentration with a Bradford assay, an equal amount of total protein extracts was incubated with ChromaTek GFP-Trap® Magnetic Agarose (Proteintech) according to the manufacturer’s instructions. The samples were eluted with boiled 2x sodium dodecyl sulfate (SDS) sample buffer, and subjected to immunoblotting analysis with anti-GFP and anti-Myc-POD antibodies.

### GFP cleavage analysis

To analyze the GFP cleavage assay, seedlings were grown on MS medium without sucrose in the light for 7 days, followed by 2-to-9-day incubation in darkness for carbon starvation stress. The harvested seedlings were weighed and homogenized with a pestle in liquid nitrogen. The 2x SDS sample buffer was added to the seedlings in proportion to their weight, and the total protein extract from each sample was boiled for 3 minutes. Subsequently, the samples were centrifuged and resolved through SDS-polyacrylamide gel electrophoresis (SDS-PAGE), followed by immunoblotting with an anti-GFP or anti-Actin antibody. Signals were detected with the SuperSignal^TM^ Femto Maximum Sensitivity Substrate or Pico PLUS Chemiluminescent Substrate.

### *Arabidopsis* carbon starvation stress

For the fixed carbon starvation treatment, seedlings were grown on MS medium without sucrose in the light for 7 days and subsequently transferred to darkness for the indicated days. After the dark treatment, the seedlings were moved to recover for one week, during which time they were photographed. To measure the protein contents of the sample, whole seedlings were harvested for the indicated days, weighed, frozen, and ground in liquid nitrogen. The ground samples were resuspended in 2x SDS sample buffer in proportion to their weight. Subsequently, the samples were centrifuged and resolved through SDS-PAGE, followed by Coomassie brilliant blue (CBB) staining and immunoblotting with an anti-RbcL or anti-Actin antibody. Signals were detected with SuperSignal^TM^ Pico PLUS Chemiluminescent Substrate.

### Real-time quantitative PCR analysis

Total RNA was prepared using the RNeasy Plant Mini Kit (QIAGEN) and reverse transcribed using SuperScript II reverse transcriptase (Invitrogen) according to the manufacturer’s instructions. Quantitative RT-PCR was performed using PowerUP^TM^ SYBRGreen Master Mix (Applied Biosystems). The primers used are listed in Supplementary Table 1. Three biological replicates were analyzed with three technical replicates per sample. The relative expression of candidate genes was normalized to *Actin2*.

### Immunoblot analysis

Three-day-old dark-grown or seven-day-old light-grown seedlings of WT and autophagy mutants were grown in MS media without sucrose, followed by treatment with ACC for 2 hours. The harvested seedlings were blotted to remove excess liquid and weighed. Subsequently, the seedlings were frozen in liquid nitrogen and ground to a fine powder. The ground samples were resuspended in 2× SDS sample buffer in proportion to their weight. An equal amount of total protein extract from each sample was boiled for 3 minutes and centrifuged to extract the total protein. The total proteins were resolved by SDS-PAGE, followed by immunoblotting with anti- CTR1 or anti-Actin antibodies. The signals were detected using SuperSignal™ Pico PLUS Chemiluminescent Substrate.

## AKOWLEDGEMENTS

This work was supported by the National Science Foundation (IOS-2245525 to GM.Y.). W.Z. was supported by the EMBRIO institute, contract #2120200, a National Science Foundation (NSF) Biology Integration Institute. We thank Dr. Taijoon Chung (Busan National University, South Korea) for providing the autophagy mutants and technical advice and Dr. Zhixiang Chen (Purdue University) for critical reading of the manuscript.

## AUTHOUR CONTRIBUTION

GMY and HLP conceived idea and designed research; HLP performed most of the experiments; WZ, YC, and CP performed autophagosome localization and quantification; GMY wrote original draft; GMY, HLP, and WZ edited the manuscript.

## CONFLITS OF INTEREST

All authors declared no conflicts of interest

## DATA AVAILABILITY

All data is available through the main text and supplementary information

