## Supplemental information for "Autophagy-mediated CTR1 turnover orchestrates the reciprocal interaction between autophagy and ethylene signaling"

**This PDF file includes**

Supplementary Figures 1-7

Supplementary Table 1

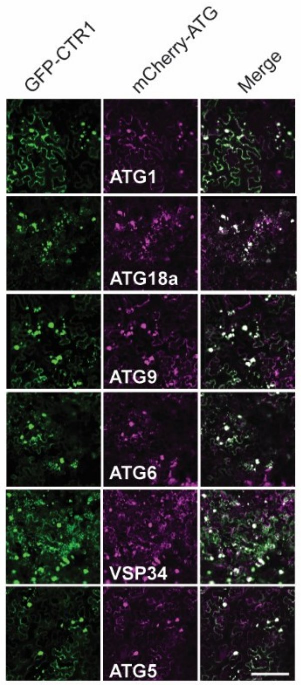


Supplementray Figure 1

**Supplementary Figure 1. CTR1 colocalizes with various ATG proteins involved in autophagosome formation.** Leaves of *N. benthamiana* were co-infiltrated with Agrobacterium carrying plasmids expressing GFP-CTR1 and the indicated mCherry-ATG proteins. After 3 days, the overlap of GFP and mCherry fluorescence was observed by confocal microscopy. Scale bar, 50 μm.

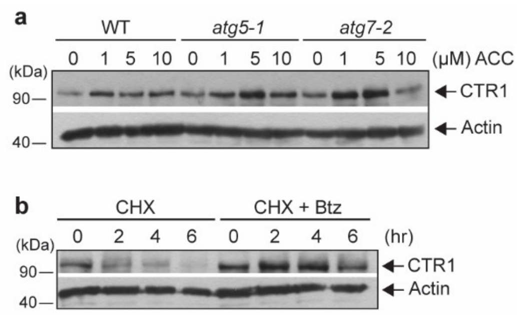


Supplementray Figure 2

**Supplementary Figure 2. ACC-induced stabilization of CTR1 is not regulated by autophagy. a**. ACC stabilizes CTR1 in both WT and *atg* mutants. Seedlings were grown in MS medium for 3 days in the dark, and treated with different concentrations of ACC in liquid MS medium. Total protein extracts were immunoblotted using anti-CTR1 and anti-Actin antibodies. Actin was detected to show even loading between samples. **b**. The stability of CTR1 is also regulated by the 26S proteasome-ubiquitin pathway. Dark-grown WT seedlings were treated with or without 50 μM Bortezomib (Btz) for 2 hrs, followed by cycloheximide (CHX) treatment to measure its degradation kinetics.

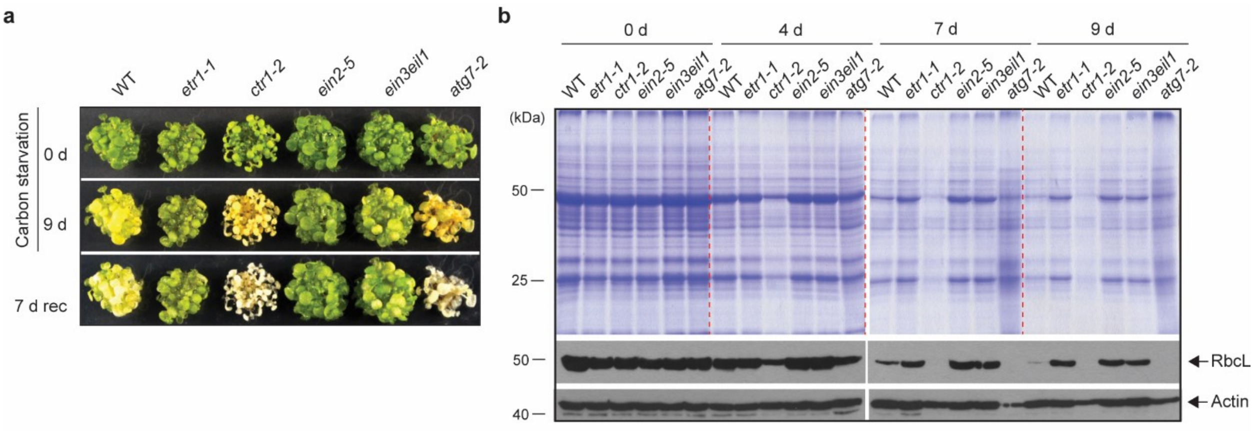


**Supplementray Figure 3**

**Supplementary Figure 3. Ethylene-insensitive mutants are hyposensitive to carbon starvation stress. a**. Carbon starvation-induced senescence phenotypes of WT, *ctr1-2*, *atg7-2*, and ethylene-insensitive mutants (*etr1-1*, *ein2-5*, *ein3eil1*). Seven-day-old light-grown seedlings were grown on MS medium without sucrose and transferred to darkness for 9 days, followed by recovery in the light for 7 days (7-rec). **b**. Coomassie brilliant blue-stained SDS-PAGE of total protein extracts of WT, *ctr1-2*, ethylene insensitive mutants, and *atg7-2* mutant. Seven-day-old light-grown seedlings were transferred to total darkness for the indicated times. Seedlings were harvested at the indicated time after carbon starvation, and total cellular proteins were resolved in SDS-PAGE, stained by CBB, or immunoblotted using antibodies against RbcL and actin (loading control).

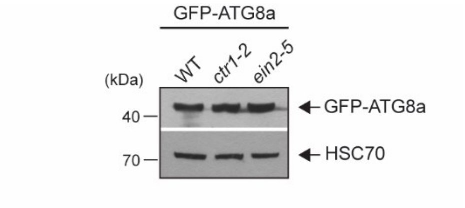


Supplementray Figure 4

**Supplementary Figure 4. Expression of GFP-ATG8a in WT, *ctr1-2*, and *ein2-5* mutants.** Similar levels of GFP-ATG8a were expressed in WT, *ctr1-2*, and *ein2-5* mutants. Total protein extracts from seedlings were immunoblotted using antibodies against GFP and HSC70 (loading control).

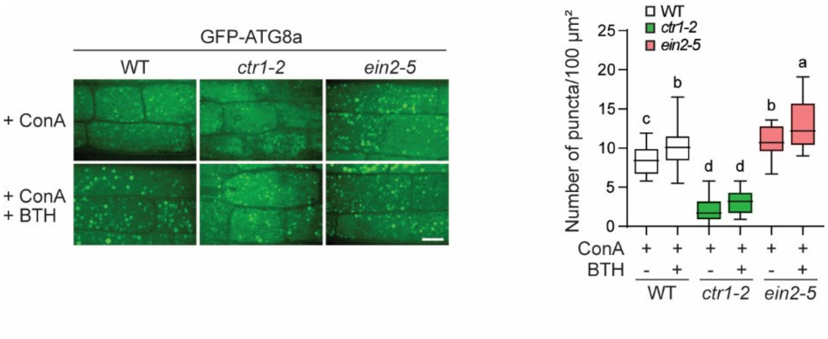


**Supplementray Figure 5**

**Supplementary Figure 5. Autophagosome formation is altered in *ctr1-2* and *ein2-5* in response to BTH treatment.** Six-day-old light-grown seedlings were transferred to sucrose-free MS medium with or without benzo-(1,2,3)-thiadiazole-7-carbothioic acid (BTH), a salicylic acid agonist. ConA was also included in both conditions to prevent vacuolar degradation of autophagosomes. Quantification of autophagosome formation in WT, *ctr1-2*, and *ein2-5*. Scale bars, 10 µm. One-way ANOVA with Tukey's post-hoc test was performed, and different letters denote statistically significant differences between groups, $p \leq 0.0001$. ($n \geq 20$, different cells from five to six seedlings for each treatment were used for measurement).

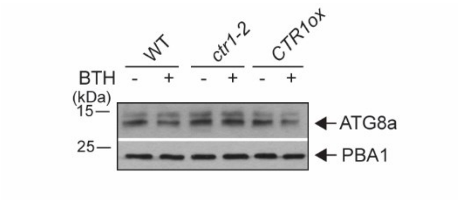


**Supplementray Figure 6**

**Supplementary Figure 6. The steady state levels of endogenous ATG8a in WT, *ctr1-2*, and CTR overexpression lines.** Four-day-old dark grown seedlings were treated with or without BTH to induce autophagy. Total protein extracts were analyzed by immunoblotting using anti-ATG8a and PBA1 (loading control) antibodies.

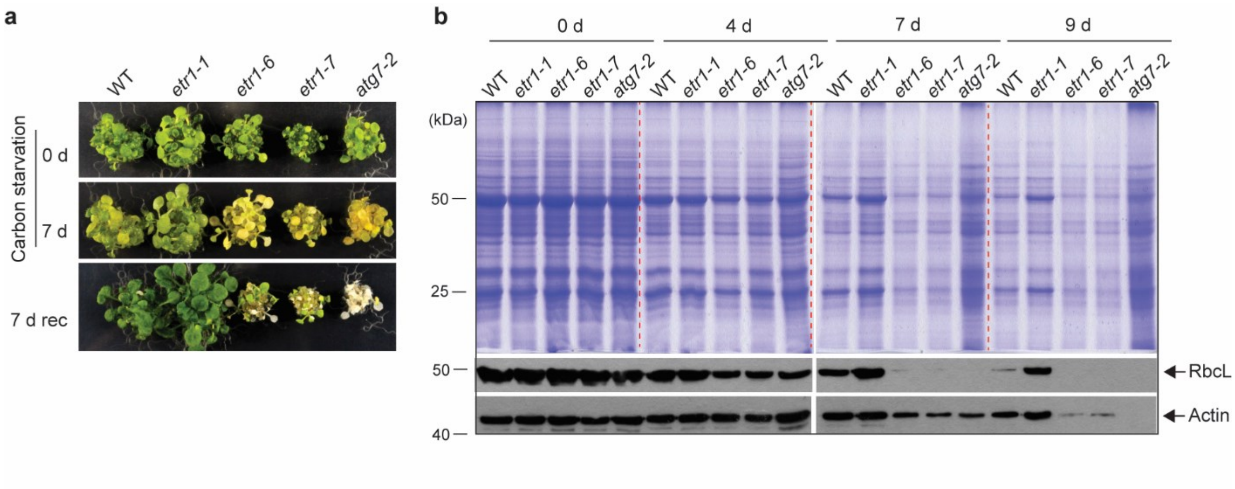


### Supplementray Figure 7

**Supplementary Figure 7. The *etr1-6* and *etr1-7* loss-of-function mutants are hypersensitive to carbon starvation stress. a**. Carbon starvation-induced senescence phenotypes of WT, *etr1-1*, *etr1-6*, and *etr1-7*. Seven-day-old light-grown seedlings were grown on MS medium without sucrose and transferred to darkness for 7 days, followed by recovery in the light for 7 days (7-rec). **b**. Coomassie brilliant blue-stained SDS-PAGE of total protein extracts of WT and *etr1* mutant seedlings. Seven-day-old light-grown seedlings were transferred to total darkness for the indicated times. Seedlings were harvested at the indicated time after carbon starvation, and total cellular proteins were resolved in SDS-PAGE, stained with Coomassie brilliant blue (CBB), or immunoblotted using antibodies against RbCL and actin (loading control).

**Supplementary Table 1. Primers used in this study**

| | |
|---|---|
| **Primers for ENTRY gateway vectors** | |
| CTR1-CACC-F | CACCATGGAAATGCCCGGTAGAAGATCT |
| CTR1-R-WS | TTACAAATCCGAGCGGTTGGGC |
| ATG8a-F | GCTTGATATCGAATTCAATGGCTAAGAGTTCCTTCAAGATC |
| ATG8a-R | CCGCTCTAGAACTAGTTCAAGCAACGGTAAGAGATCCAA |
| ATG8a-F for mCherry | ACGGATCCATGGCTAAGAGTTCCTTCAAGA |
| ATG8a-R for mCherry | TGGCGGCCGCTCTAGATCAAGCAACGGTAAGAGATCC |
| ATG1-F | GGGGCCCCCCTCGAGATGGAGTCGGCACGACTTGT |
| ATG1-R | TGCTCACCATGGATCCAAGATGAGACCGACGATGCTGT |
| ATG5-F | GGGGCCCCCCTCGAGATGGCGAAGGAAGCGGTC |
| ATG5-R | TGCTCACCATGGATCCCCTTTGAGGAGCTTTCACAAGGAC |
| ATG6-F | GGAGGCCAGTGAATTCATGAGGAAAGAGGAGATTCCAG |
| ATG6-R | CGAGCTCGATGGATCCCTAAGTTTTTTACATGAAGGCTT |
| ATG9-F | GGGGCCCCCCTCGAGATGATGAGCAGTGGGCATAAGGG |
| ATG9-R | TGCTCACCATGGATCCCCGTAATGTGGTGCTTGATGTTG |
| ATG18A-F | GGGGCCCCCCTCGAGATGGCCACCGTATCTTCTTCCTC |
| ATG18A-R | TGCTCACCATGGATCCGAAAACTGAAGGCGGTTTCAGACAG |
| VPS34-F | GGGGCCCCCCTCGAGATGGGTGCGAACGAGTTTC |
| VPS34-R | TGCTCACCATGGATCCACGCCAGTATTGAGCCCATCTG |
| CIP8-F | GGGCCCCCCTCGAGATGTCCGATGCTCCGTCGTCTTC |
| CIP8-R | CGCTCTAGAACTAGTGTAACGAGAAGTTGAAGAAGAAGAAG |
| **Primers for quantitative RT-PCR** | |
| ERF1-RT-F | ACGTTCTCAACCGCCTACAG |
| ERF1-RT-R | CGGACTCGCTCTCTGGTG |
| CTR1-RT-F | CATGGAAGCGTCCATCATTTGCA |
| CTR1-RT-R | TGGGCAGCAAAAGAATGCTGAG |
| ATG1a-RT-F | TGAACCCAGATCCAACCACG |
| ATG1a-RT-R | GAGTGGCAGCACTTGTTTCG |
| ATG2-RT-F | AATGGATAGCAAGTGGAAGC |
| ATG2-RT-R | AGATAGACCTACCGTTAGCC |
| ATG5-RT-F | GAAGGAAGCGGTCAAGTATGT |
| ATG5-RT-R | TCTTGGTGCTAACACAAGAGC |
| ATG7-RT-F | GAAGATTGTCTAGGTCGTGG |
| ATG7-RT-R | CCTGCTTTCTCTTGTATCGG |
| ATG8a-RT-F | GACAATTTGTATACGTGGTTCGT |
| ATG8a-RT-R | TCAAGCAACGGTAAGAGATCCA |
| ATG9-RT-F | GGATGATGTCCGCTTGGAGT |
| ATG9-RT-R | TCTCTGCTCACGACGAGTTG |
| ATG11-RT-F | ACCGGGAGGAATTATTGGAG |
| ATG11-RT-R | GGACTTCAAGACGACCGAAT |
| ATG13a-RT-F | ATTTCAGCTCGGAGTGGTCG |
| ATG13a-RT-R | GGCATCTGTAGAAGCTCCCAT |
| ATG18a-RT-F | CAAAGGAGTGTTACCGAGGTAT |

| | |
|---|---|
| ATG18a-RT-R | ATCCATGCCAAGAATAACAACG |
| TOR1-RT-F | ATACGCTGCCTGTGGGAAAT |
| TOR1-RT-R | CGGACAGCCAAAAGTGCTTG |
| RAPTOR1B-RT-F | GCGGTCCGCAAACTGTAATC |
| RAPTOR1B-RT-R | TGGAACTCTGTGCTTGGGTC |
| LST8-1-RT-F | TCCTGCAAACAAATATCTAGCGAC |
| LST8-1-RT-R | CCATCCACTGAGAAGACGCA |
| ACTIN2-RT-F | ATTCAGATGCCCAGAAGTCTTGTT |
| ACTIN2-RT-R | ACGGTCAGCGATACCTGAGAAC |
